# Examining Reproducibility of EEG Schizophrenia Biomarkers Across Explainable Machine Learning Models

**DOI:** 10.1101/2022.08.16.504159

**Authors:** Charles A. Ellis, Abhinav Sattiraju, Robyn Miller, Vince Calhoun

## Abstract

Schizophrenia (SZ) is a neuropsychiatric disorder that adversely effects millions of individuals globally. Current diagnostic efforts are symptom based and hampered due to the variability in symptom presentation across individuals and overlap of symptoms with other neuropsychiatric disorders. This spawns the need for (1) biomarkers to aid with empirical SZ diagnosis and (2) the development of automated diagnostic approaches that will eventually serve in a clinical decision support role. In this study, we train random forest (RF) and support vector machine (SVM) models to differentiate between individuals with schizophrenia and healthy controls using spectral features extracted from resting state EEG data. We then perform two explainability analyses to gain insight into key frequency bands and channels. In our explainability analyses, we examine the reproducibility of SZ biomarkers across models with the goal of identifying those that have potential clinical implications. Our model performance results are well above chance level indicating the broader utility of spectral information for SZ diagnosis. Additionally, we find that the RF prioritizes the upper γ-band and is robust to loss of information from individual electrodes, while the SVM prioritizes the α and θ-bands and P4 and T8 electrodes. It is our hope that our findings will inform future efforts towards the empirical diagnosis of SZ and towards the development of clinical decision support systems for SZ diagnosis.

## I. Introduction

Schizophrenia (SZ) affects approximately 20 million people globally [1]. Because symptoms of SZ overlap with other neuropsychiatric disorders and can vary from individual to individual, it can be difficult to diagnose and, in turn, to treat [1], [2]. As such, there is a need for automated diagnostic methods and for the identification of novel biomarkers. In this study, we use several explainable machine learning methods for automated schizophrenia diagnosis and to identify biomarkers of schizophrenia that are reproducibly important across models.

Diagnosing schizophrenia is challenging, as it is typically diagnosed based upon symptoms rather than any specific neurological or electrophysiological biomarkers [2]. Its symptoms can include lost cognitive function, delusions, hallucinations, loss of pleasure, and a flattening of emotion [1]. Additionally, its symptoms can overlap with other neuropsychiatric disorders, like bipolar disorder [2], and can vary from person to person. This difficulty has led to extensive research efforts to (1) identify biomarkers than can enable the empirical SZ diagnosis and (2) develop automated SZ diagnosis approaches for clinical decision support.

Much of the research focused on SZ diagnosis and biomarker discovery has involved electroencephalography (EEG), though efforts have been made with other modalities like magnetoencephalography (MEG) [3] and functional magnetic resonance imaging (fMRI) [4], [5]. EEG offers advantages over other modalities in that it is relatively cheap and easy to collect while also having high temporal resolution [6].

Historically, many studies have developed automated SZ diagnosis approaches using EEG. These studies have involved both machine learning (ML) [7], [8] and deep learning (DL) methods [1], [9]. ML and DL each have their respective advantages and disadvantages. Traditional ML studies use extracted features, like spectral power [7], [8] or inter-channel connectivity [7], [10], and offer enhanced explainability. Spectral features do offer some advantages over connectivity regarding explainability. Namely, key EEG frequency bands can be identified through visual inspection in a clinical setting, increasing the likelihood that any identified features will have an eventual clinical impact. Although substantial progress has recently been made in DL explainability for raw EEG classification [11], [12], that is still a challenge for the space, and to our knowledge, only one deep learning study for SZ diagnosis has used explainability [9]. As such, for the goal of biomarker identification, simpler ML methods should be used where possible and to the extent that they enable accurate diagnosis.

There are some gaps in the current landscape of traditional machine learning studies involving automated SZ diagnosis. For example, many studies do not have strong cross-validation approaches or do not clearly articulate their approaches [8], [10]. By distributing segments from the same recordings across training, validation, and test groups, they artificially amplify their model performance and do not test the generalizability of the patterns learned by their models. This is not the case for all studies in the field [7], [13]. Additionally, explainable machine learning studies have typically not compared the reproducibility of explanations from multiple model types.

As such, in this study, we train a pair of explainable machine learning models on spectral features extracted from raw EEG. We then compare the explanations across models, gaining insight into the relative importance of each of the frequency bands and channels to the models, and compare the explanations with existing literature. Our study, to the best of our knowledge, represents the first study of resting state EEG-based classification of SZ with spectral features that has sought to understand the reproducibility of identified SZ biomarkers via a comparison across explainable machine learning methods.

## II. Methods

In this section, we describe and discuss our analysis approach. An overview of our approach is shown in Figure 1.

**Figure 1.**
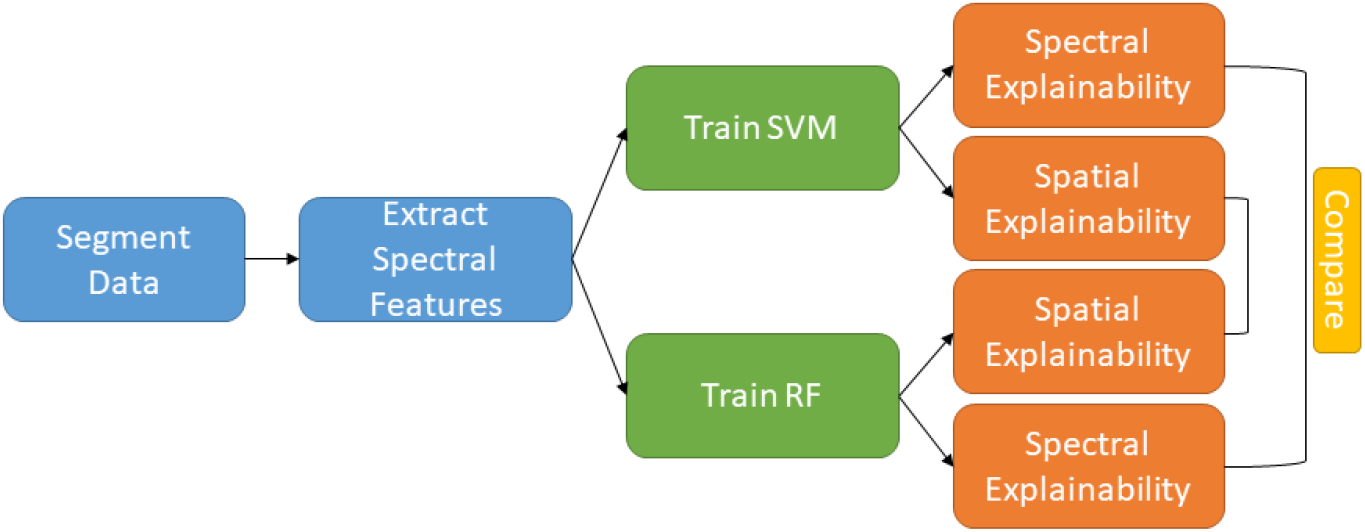
Methods Flow Chart. Our analysis consisted of several stages: preprocessing, model development, explainability, and comparison of explanations. The preprocessing stage (i.e., blue boxes) mainly consisted of data segmentation and spectral feature extraction. In the model development stage (i.e., green boxes), we trained and tested an SVM and RF. In our explainability analysis (i.e., orange boxes), we examined the importance of each frequency band and channel to the model performance. Lastly, we compared the explanations across models (i.e., gold box).

### A. Data Collection

We utilized a publicly available scalp EEG dataset [14] that has been used in previous studies [2]. The dataset contains 14 individuals with SZ (SZs) and 14 healthy controls (HCs), collected at the Institute of Psychiatry and Neurology in Warsaw, Poland. All participants gave informed consent prior to the study. Recordings were performed in a 15-minute resting state at 250 Hz. During recording, 64 electrodes were placed in the standard 10-20 format.

### B. Data Preprocessing

Like previous studies [9], we only analyzed the Fp1, Fp2, F7, F3, Fz, F4, F8, T3, C3, Cz, C4, T4, T5, P3, Pz, P4, P6, T6, O1, and O2 electrodes. To maximize the amount of data available for training, recordings were separated into 25-second epochs using a sliding window approach with a step size of 2.5s. We then randomly upsampled HC samples to ensure balanced representation of both classes. After segmenting the data, we converted each sample to the frequency domain with a Fast Fourier Transform (FFT). For each electrode, we calculated the average phase and amplitude within each frequency band (i.e. δ (0 – 4 Hz), θ (4 - 8 Hz), α (8 – 12 Hz), β (12 – 25 Hz), γ_lower_ (25 – 55 Hz), and γ_upper_ (65 – 125 Hz)) for all individuals. We then z-scored each feature individually across all samples.

### C. Model Development

We developed a Random Forest (RF) and Support Vector Machine (SVM) in this study. We trained both models using a nested cross-validation strategy with 10 splits of 80% training-validation data and 20% hold-out testing data. Within each inner fold, the training data was split into 10 folds of 64% training data and 16% validation data. Data was split by subject rather than by sample, ensuring the model would evaluate complete subject data in the test set that were unseen in the training set. We tuned both models’ hyperparameters on the training and validation splits, and due to problems with overfitting, the best hyperparameters for each model were selected from the model with minimal absolute difference in mean validation sensitivity (SENS) and specificity (SPEC). After selecting the optimal hyperparameters, each model was refit using the training and validation data from its respective fold. When testing the model, we calculated the mean (μ) and standard deviation (σ) of the F1 score, accuracy (ACC), SENS, and SPEC.

### D. Model Explainability

For insight into the importance of the features to the models and to help identify biomarkers, we used a permutation feature importance approach [15] with a grouping mechanism that enabled us to perform two separate analyses. (1) We permuted the features associated with each canonical frequency band for spectral importance. (2) We permuted the features associated with each channel for spatial importance. Following each permutation, we calculated the percent change in the test ACC. When permuting, all feature values in a group for a particular subject were assigned to one other randomly selected subject. We performed 100 permutations for each analysis.

## III. Results and Discussion

In this section, we describe and discuss the results of our automated SZ diagnosis approach and the biomarker discovery efforts of our explainability approach.

### A. Automated Diagnosis of Schizophrenia

Table 1 shows the μ and σ of the performance results of each of our models. Both classifiers performed above chance-level, with the RF outperforming the SVM in all metrics except SENS. The mean SPEC was much higher for the RF than the SVM, indicating that the RF had more success correctly classifying HC instances. Our SVM accuracy was lower than past studies that have applied SVMs to various spectral features to detect SZ [16], [17]. This is likely due to our more robust subject-based cross-validation approach and choice of spectral features [16] or use of resting state, rather than task, EEG data [17]. Another past study that used an RF to train on amplitude data of over 200 frequency widths performed at an accuracy 20% higher than the RF in this study; however, it utilized a cross-validation approach that allowed samples from the same subjects to be assigned to training, validation, and testing sets simultaneously, which would inflate model performance [8]. In unreported instances in which we used a similar cross-validation approach to [8] to train our SVM and RF, we saw similarly high levels of performance.

**TABLE I.** Model Performance Results

|  | F1 | ACC | SENS | SPEC |
| --- | --- | --- | --- | --- |
| **SVM** | 68.88 ± 12.85 | 70.60 ± 05.60 | 77.65 ± 18.08 | 59.99 ± 19.88 |
| **RF** | 72.54 ± 09.85 | 74.99 ± 08.19 | 75.46 ± 15.45 | 77.87 ± 11.82 |

### B. Schizophrenia Biomarkers Identification

Figure 2 shows our spectral explainability results for both models. The SVM prioritized θ and α most, while the RF relied heavily on γ_upper_. Excluding γ_upper_, the RF was more robust to the permutation of features associated with various frequency bands than the SVM. The SVM results support past findings on the difference in θ and α power between HCs and SZs [18]. The results for the RF also corroborate past findings of the distinction between HCs and SZs in the γ-bands [2]. The differences in explanations between the two classifiers indicate that there are multiple ways to diagnose SZ based upon resting state EEG spectral features and hint at their potential utility as biomarkers.

**Figure 2.**
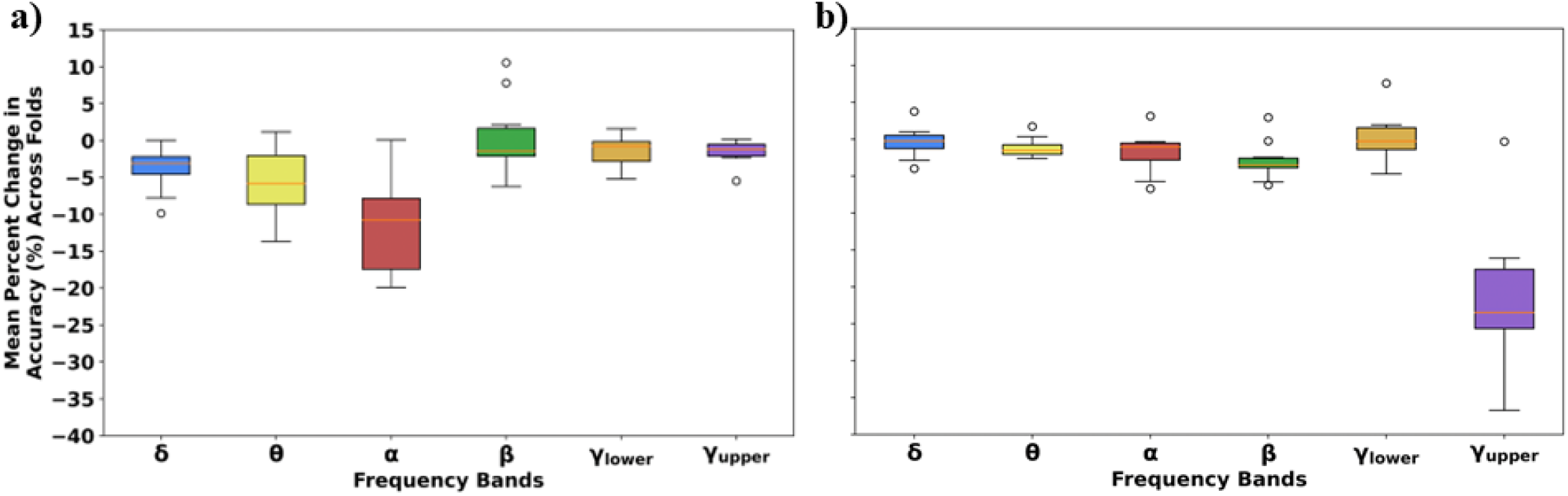
Spectral Importance Results. Panel a) and b) show the importance of each frequency for the SVM and RF, respectively. The frequency bands are arranged on the x-axis. The y-axes are aligned across panels, showing the mean percent change in accuracy across all 100 permutations within each fold. Note that for the SVM, the α-band is most important. In contrast, for the RF, the γupper band is most important.

Figure 3 shows our spatial explainability results for both models. The RF was much more robust to the permutation of features associated with each channel than the SVM, which found the P4, T8, and P3 channels to be the most important for SZ classification. These electrodes are in temporal and parietal regions of the brain, which are associated with the development of SZ [19]. The differences in features learned between models could possibly be explained by the difference in model structure or a correlation between many of the channel-related features, which would decrease the importance attributed to each feature.

**Figure 3.**
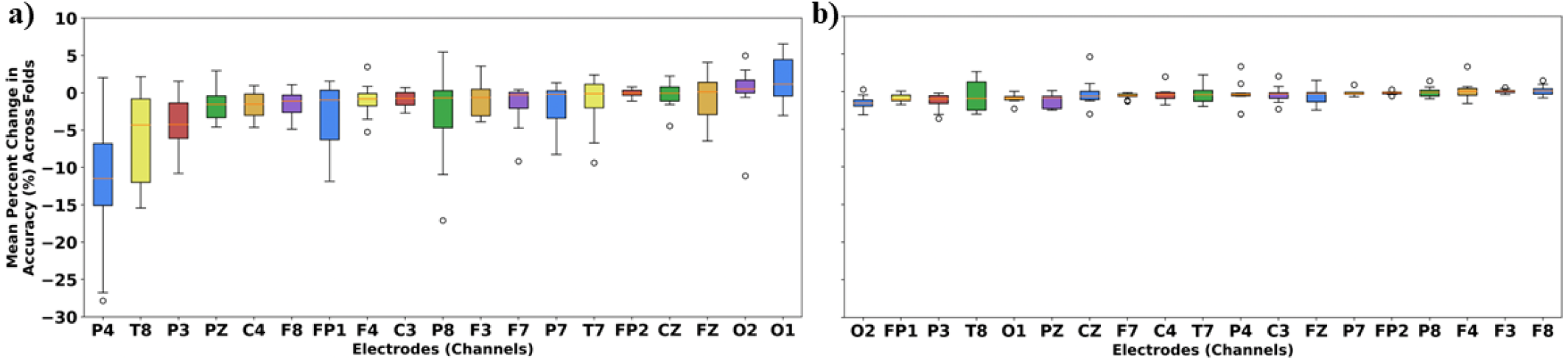
Channel Importance Results. Panel a) and b) show the importance of each channel for the SVM and RF, respectively. The electrodes are arranged from left to right on the x-axis in order of the magnitude of their importance. The y-axes are aligned across panels, showing the mean percent change in accuracy across all 100 permutations within each fold. Note that the RF was much more robust to the loss of any individual channel. In contrast, the SVM was reliant upon the P4 and T8 electrodes for its performance.

### C. Limitations and Next Steps

We included both phase and amplitude information in our analysis. In the future, it would be beneficial to include and explainability analysis examining the relative importance of the phase features relative to the amplitude features. Additionally, in our analysis, we included two machine learning classifiers capable of identifying nonlinear patterns. In a future analysis, it would be interesting to compare their performance to a simpler linear model like logistic regression. Additionally, while we compared the reproducibility of our findings across models, it would also be interesting to compare the reproducibility of our results across datasets. It would also be interesting to examine the interactions between features that might have been found by the models using explainability approaches like SHAP.

## IV. Conclusion

In this study, we trained two machine learning models for automated schizophrenia diagnosis with a robust cross-validation approach that appropriately assessed model generalizability. We then implemented and compared a set of explainability analyses for each model that provided insight into the effects of schizophrenia upon resting state EEG activity. The features that we identified as important for each classifiers fit well with existing literature and help build up the reliability of said biomarkers for use in a clinical setting. Additionally, the differences in explainability results, paired with our robust cross-validation approach, highlight the richness of the EEG features that we examined and demonstrated that there are multiple distinct groups of spectral EEG features that can contribute to SZ diagnosis. It is our hope that this work will inform future efforts towards automated SZ and neuropsychiatric disorder diagnosis and also contribute to the eventual empirical diagnosis of schizophrenia.

